# Nascent evolution of recombination rate differences as a consequence of chromosomal rearrangements

**DOI:** 10.1101/2023.03.27.534312

**Authors:** K. Näsvall, J. Boman, L. Höök, R. Vila, C. Wiklund, N. Backström

**Affiliations:** Evolutionary Biology Program, Department of Ecology and Genetics, Uppsala University. Norbyvägen 18D, 752 36 Uppsala, Sweden; Butterfly Diversity and Evolution Lab, Institut de Biologia Evolutiva (CSIC-UPF), Barcelona, Spain; Department of Zoology: Division of Ecology, Stockholm University, Stockholm, Sweden

## Abstract

Reshuffling of genetic variation occurs both by independent assortment of chromosomes and by homologous recombination. Such reshuffling can generate novel allele combinations and break linkage between advantageous and deleterious variants which increases both the potential and the efficacy of natural selection. Here we used high-density linkage maps to characterize global and regional recombination rate variation in two populations of the wood white butterfly (*Leptidea sinapis*) with distinct karyotypes. The recombination data were compared to estimates of genetic diversity and measures of selection to assess the relationship between chromosomal rearrangements, crossing over, maintenance of genetic diversity and adaptation. Our data show that the recombination rate is influenced by both chromosome size and number, but that the difference in recombination rate between karyotypes is reduced as a consequence of a higher frequency of double crossovers in larger chromosomes. As expected from effects of selection on linked sites, we observed an overall positive association between recombination rate and genetic diversity in both populations. Our results also revealed a significant effect of chromosomal rearrangements on the rate of intergenic diversity change between populations, but limited effects on polymorphisms in coding sequence. We conclude that chromosomal rearrangements can have considerable effects on the recombination landscape and consequently influence both maintenance of genetic diversity and efficiency of selection in natural populations.

**Author summary:** Reshuffling genetic variation is fundamental for maintaining genetic diversity and creating novel allelic combinations. The two main processes involved are the independent assortment of chromosomes and homologous recombination. The number and size of chromosomes can influence the amount of pairwise reshuffling and local recombination patterns. However, studying this in natural populations is challenging. In this study, we used the wood white butterfly, which exhibits an extreme within-species karyotype difference. Extensive fusions and fissions have resulted in almost twice as many chromosomes in the southern populations compared to the northeast populations. This unique system allowed us to assess the relationship between karyotype differences, pairwise reshuffling, recombination rate variation and subsequent effects on diversity and linked selection. We found that a higher number of chromosomes result in a higher recombination rate, although the difference was less than expected due to multiple recombination events occuring on longer chromosomes. Both populations showed an association between recombination rate and genome-wide patterns of genetic diversity and efficacy of selection. We provide evidence that chromosomal rearrangements have considerable effects on the recombination landscape and thereby influence the maintenance of genetic diversity in populations.

## Introduction

Genetic variation is a prerequisite for evolutionary change and mutation is the ultimate source of novel genetic variants. However, mutation only contributes to variation at individual genomic sites and establishment of new combinations of alleles across loci is dependent on reshuffling via independent segregation and recombination. Physical linkage can lead to a reduction in genetic diversity as a consequence of fixation or loss of haplotype blocks, either by random drift, or more rapidly by selection via hitch-hiking and/or background selection [1,2]. Recombination events resolved as crossovers (from here on recombination) breaks physical linkage and generally leads to maintenance of genetic variation by generating a higher number of segregating haplotypes in the population [3–5]. Physical linkage between variants influenced by natural selection can also result in less efficient selection than if such variants would have been segregating independently. This process is generally referred to as Hill-Robertson interference, and can lead to an increased accumulation rate of deleterious mutations in regions with reduced recombination rate [6,7]. However, reshuffling can also break up co-adapted or synergistically interacting loci and therefore hinder beneficial epistasis.

Reshuffling of genetic variation occurs by two different mechanisms in diploid organisms. First, the independent assortment of homologous chromosomes to the gametes during meiosis (Mendel’s second law) results in reshuffling of loci located on different chromosomes. Hence, the higher the number of chromosomes in an organism, the higher the potential for pairwise reshuffling. However, the effect renders a diminishing return as the probability of pairwise reshuffling approaches the maximum of 0.5 [8]. Second, the exchange of genetic material between chromosome pairs during homologous recombination can lead to novel allele combinations within chromosomes. Consequently, characterizing where and how often recombination events occur along chromosomes is key to understanding maintenance of genetic diversity and the efficiency of selection in different parts of the genome. Empirical data have unveiled that the recombination rate differs over several orders of magnitude, both among species, between chromosomes and across different genomic regions [9–11]. Interspecific variation in genome wide recombination frequency could be related to differences in haploid chromosome number [10] since at least one recombination event per chromosome pair is necessary for correct segregation during meiosis in many organisms [9–11]. However, there are exceptions to this generalisation. In Lepidoptera females and some Diptera males, for example, where meiotic divisions are achiasmatic [12–14]. The location and frequency of crossover events along chromosomes has been investigated in several different organisms. The results so far generally show that the recombination landscape can be highly heterogeneous and that location and frequency of recombination events can be affected by, for example, interference between chiasmata, presence of centromeres and telomeres, nucleotide composition and chromatin state [9,10,15–17]. However, despite the potential importance of chromosomal rearrangements on the recombination landscape and recombination-dependent evolutionary processes, detailed analyses of links between karyotype changes and recombination are limited, in particular in natural populations where fine-scale recombination rate data have been difficult to establish.

In order to explore how karyotype differences affect the recombination rate in a natural system, we used the wood white butterfly (*Leptidea sinapis*), a species within the Eurasian genus *Leptidea* which have considerable interspecific karyotype variability compared to the majority of other lepidopterans which usually have a remarkably stable karyotype of 2n ∼ 62 [18]. Within *L. sinapis*, there is also extreme intraspecific variation in karyotype; chromosome numbers range from 2n = 56 - 62 in the northern (Scandinavia) and eastern (central Asia) parts to 2n = 108 - 110 in the south-western (Iberian peninsula) part of the distribution range [19,20]. We know from genomic data that the intraspecific karyotype rearrangements in *L. sinapis* predominantly have been driven by recurrent fissions and fusions and that fission/fusion polymorphisms currently segregate in different populations [21]. Previous analyses in other butterfly species have unveiled an inverse relationship between chromosome size and recombination rate [22], and a positive association between chromosome count and neutral diversity across Lepidoptera [23,24]. In addition, a detailed analysis in *Heliconius ssp.* has shown that fused chromosomes have a reduced recombination rate and genetic diversity as compared to chromosomes not involved in rearrangements [25,26]. Here, we use a study system that allows us to investigate the effects of extensive chromosomal rearrangements, both fissions and fusions, on the recombination landscape and recombination-dependent evolutionary processes.

The aims of the study were; i) to characterize the recombination landscape in Swedish (2n = 56, 57) and Catalan (2n = 106 - 110) populations of *L. sinapis*, two populations with highly different karyotypes, ii) to compare the genome-wide probability of pairwise reshuffling between the two populations, and, iii) to quantify the effect of chromosomal rearrangements on maintenance/loss of genetic diversity and the efficacy of selection. We hypothesized that the Catalan population, with a more fragmented karyotype, would have a higher recombination rate and a higher total pairwise reshuffling rate, resulting in reduced linkage disequilibrium, higher neutral genetic diversity and increased efficacy in removal of slightly deleterious mutations.

## Results

### Recombination rates

We used pedigree data from a previous study to estimate the recombination rate from linkage maps for the two populations. The total recombination distance was considerably longer for the Catalan (2,300 cM) than for the Swedish (1,711 cM) population (Suppl. Fig. 1, Suppl. Table 1). Despite a shorter recombination distance per chromosome in the Catalan (average = 43 cM; range = 10 - 90 cM) compared to the Swedish population (average = 59 cM; range 33 - 84 cM), the average recombination rate in the Catalan population was significantly higher (mean and standard deviation; 4.25 ± 211 cM / Mb) than in the Swedish population (3.03 ± 641 cM/Mb) (Wilcoxon’s test; W = 1,726,620, p-value = 4.33*10^-02^; Fig. 1). In the Catalan population, the recombination rate was lower on the Z-chromosome than on the autosomes, but this difference was not significant (Z-chromosomes = 2.86 ± 139; Autosomes = 4.46 ± 217 cM / Mb; Wilcoxon’s test; W = 168,928; p-value = 3.99*10^-01^). This effect can be a consequence of the relative difference in size between the Z-chromosomes and the autosomes in the two different populations. The Z-chromosomes are virtually conserved between the two populations and hence constitute the largest chromosomes in the Catalan population, except for one large autosome. In the Swedish population on the other hand, the sizes of the Z-chromosomes are comparable to the majority of autosomes and the recombination rates of the two chromosome classes were also very similar (Z = 3.03 ± 36.3; A = 3.03 ± 675 cM / Mb).

**Fig. 1.**
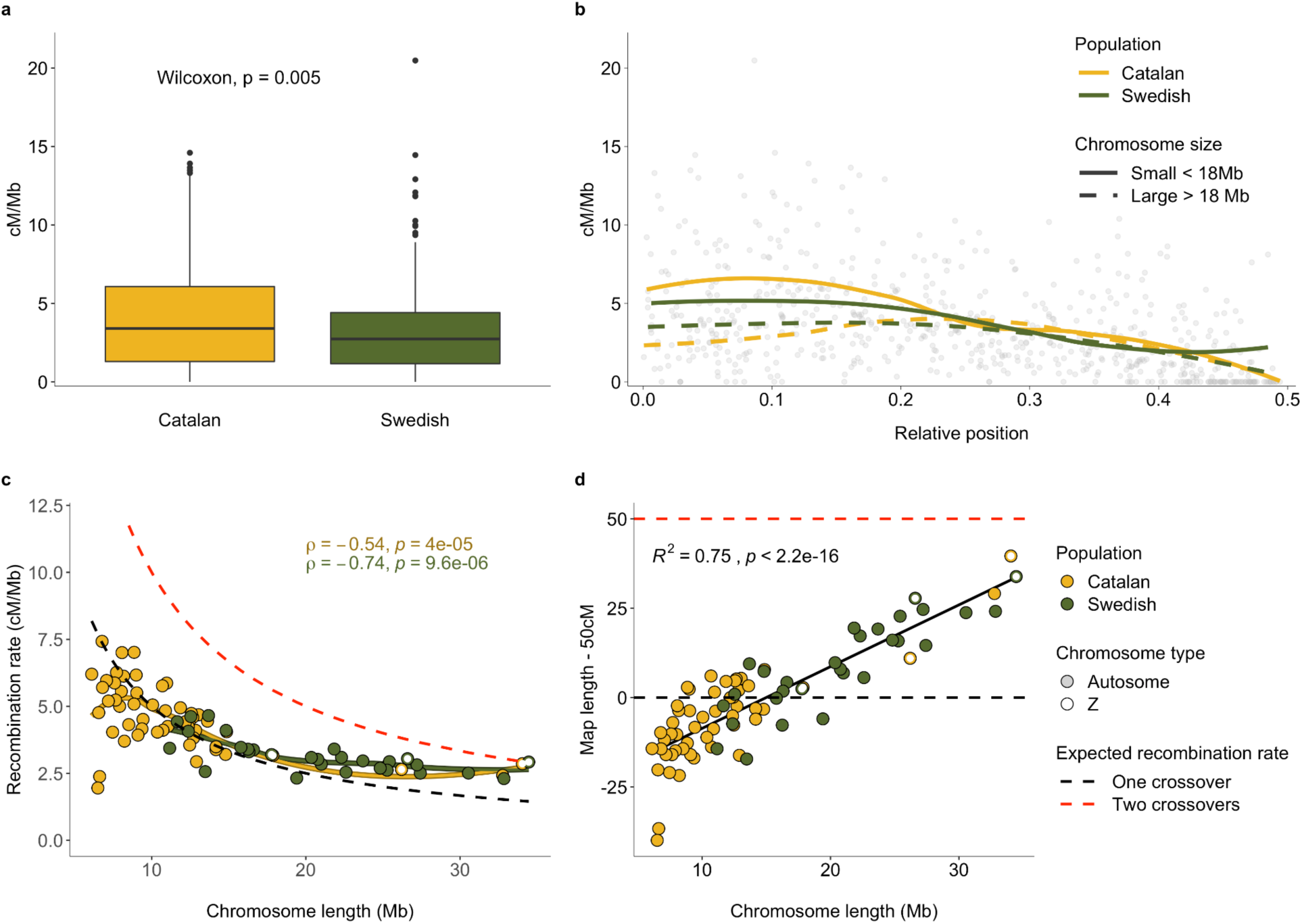
a) The distribution of recombination rate estimates (cM, y-axis) in the Catalan and Swedish populations, respectively. Box hinges represent the 25th and 75th percentiles, whiskers extend to values within 1.5 times the distance between the 25th and 75th percentiles and solid dots represent outliers. b) Regional distribution of the recombination rate (cM / Mb; y-axis) for relative positions from the center (0) to the end (0.5) of chromosomes in the Swedish (green) and the Catalan (yellow) populations, respectively. Lines represent local regression (LOESS) of large (> 18 Mb; dashed lines) and small chromosomes (< 18 Mb; solid lines), respectively. c) Association between the weighted mean recombination rate (cM / Mb; y-axis) and chromosome size (Mb; x-axis) in the Swedish (green) and Catalan (yellow) populations. The dashed lines represent the expected recombination rate with one (black) and two (red) crossovers per meiosis. The Z-chromosomes are represented by open and the autosomes by filled circles. d) Excess map length (map length −50 cM; y-axis) as a function of chromosome size. Colors and symbols as in c) and the regression line and statistics correspond to a linear regression model.

We observed a significant negative association between the recombination rate and chromosome size in both the Swedish (Spearman’s rank correlation, rho = −0.51, p-value = 5.1*10^-3^) and the Catalan population (rho = −0.44, p-value = 1.2*10^-3^; Fig. 1). The larger chromosomes in the Catalan population had similar recombination rates as chromosomes of comparable size in the Swedish population, suggesting that the overall difference in total linkage map length and average recombination rate between the populations is predominantly an effect of the difference in karyotype structure. We also found that the recombination rate on smaller chromosomes was close to the expected rate (50 cM / chromosome) with a single crossover per meiosis (Fig. 1). However, the recombination rate on the larger chromosomes was higher, showing that more than one crossover per meiosis can occur if a chromosome is large enough (> ∼ 18 Mb). The ‘excess map length’ (remaining after removing the assumed obligate crossover (50 cM) from chromosome specific estimates) was significantly positively associated with chromosome size and explained 75% of the variation (linear regression model, R^2^ = 0.75, p-value = 2.2*10^-16^; Fig. 1).

The regional distribution of recombination events showed a bimodal pattern for the larger chromosomes (> ∼ 18 Mb) with a pronounced drop in the center and at the ends (Fig. 1, Suppl. Fig. 2). For smaller chromosomes (< ∼ 18 Mb) on the other hand, the recombination rate was highest in the center (Fig. 1, Suppl. Fig. 2). The decrease in recombination in the chromosome ends was less pronounced in the small chromosomes in the Swedish population (Fig. 1). This could be a consequence of that several of the smaller chromosomes are involved in ‘fusion polymorphisms’ currently segregating in the Swedish population.

### Pairwise reshuffling

We proceeded by estimating the relative contribution of independent segregation and homologous recombination to the total reshuffling rate in the different populations. The major mechanism for genome-wide reshuffling was the number of chromosomes in both populations (Table 1). The male interchromosomal contribution was 106 times higher than the intrachromosomal contribution in the Catalan and 61 times higher in the Swedish population. Despite the lower recombination rate on larger chromosomes, the contribution of intrachromosomal pair-wise reshuffling increased linearly with chromosome size and size explained 62% of the variation between chromosomes (Fig. 2). This is likely a consequence of the occasional occurrence of double crossovers on larger chromosomes which should increase the pairwise reshuffling rate.

**Table 1.** Estimates of the male intrachromosomal (Intra male) and interchromosomal (Inter male) contribution to genome-wide reshuffling. The total contribution of males (Total male) and females (Inter female) and sex-average reshuffling (Sex average) are also given for comparison.

| Population | Intra male | Inter male | Total male | Intra female | Inter female | Sex average |
| --- | --- | --- | --- | --- | --- | --- |
| Catalan | 0.0045 | 0.488 | 0.493 | 0 | 0.482 | 0.487 |
| Swedish | 0.0079 | 0.481 | 0.489 | 0 | 0.476 | 0.483 |

**Fig. 2.**
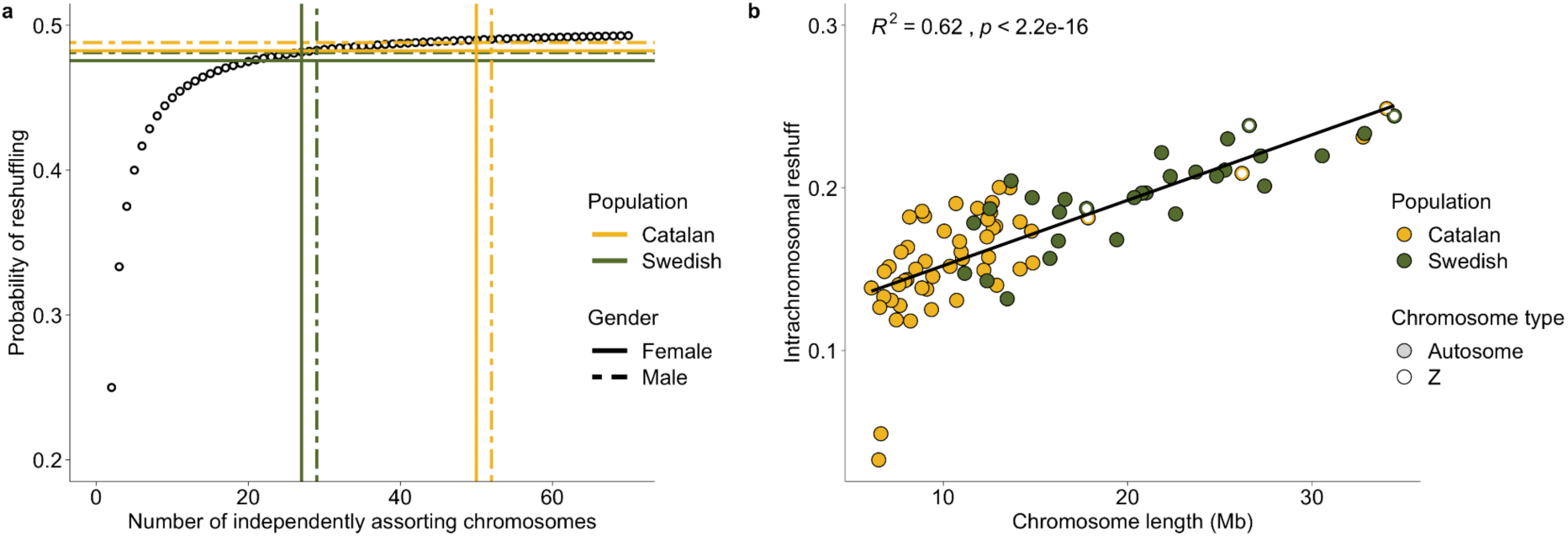
a) Genome-wide probability of pairwise reshuffling per population and sex as a function of number of independently segregating chromosomes (interchromosomal effect). The open circles represent the expected trajectory of reshuffling in genomes with equal-sized chromosomes as the chromosome number increases (x-axis). b) The probability of pairwise reshuffling within each chromosome (y-axis) as a function of chromosome size (x-axis) in the two populations (intrachromosomal reshuffling). The line represents a linear regression model.

The analysis revealed that the proportional difference in total reshuffling is less than 1% higher in the Catalan than in the Swedish population, despite the fact that the difference in chromosome numbers is almost two-fold and that females contribute more to the total reshuffling rate in the Catalan population (Table 1). The difference in reshuffling between populations was further reduced by the higher probability of intrachromosomal reshuffling in the Swedish males - the genome-wide intrachromosomal reshuffling contribution was 75.6% higher in the Swedish population (Table 1).

### Associations between recombination and genetic diversity

Given the observed variation between populations in both overall map length, average recombination rate and relative intrachromosomal reshuffling rate, we proceeded by investigating the relationship between recombination rate and genetic diversity. We found that the Catalan population had significantly higher *π* in all site categories (Wilcoxon’s test; p-values = 2.73*10^-04^- 7.05*10^-07^), but we did not detect any difference between the populations in the ratio of zero- (*π*_0_) to four-fold (*π*_4_) degenerate sites (p-value = 1.52*10^-01^; Table 2). Consistent with the ongoing and dynamic karyotype changes in these populations, there was also no significant association between *π* per chromosome and chromosome length after excluding the Z-chromosomes (Suppl. Fig. 3). However, the regional distribution of *π* estimates followed the expectations from the nearly neutral theory in both populations, with a decrease in *π* and an increase in *π*_0_ / *π*_4_ at the terminal ends of the chromosomes where the recombination rate is significantly reduced (Fig. 3). We also observed a lower neutral genetic diversity for the Z-chromosomes compared to the autosomes. In addition, while Z1 and Z3 had similar *π*_0_ / *π*_4_-ratios as the autosomes, Z2 had a significantly higher *π*_0_ / *π*_4_-ratio compared to the other chromosomes in both populations, driven by an elevated *π*_0_ (Suppl. Fig. 3).

**Fig. 3.**
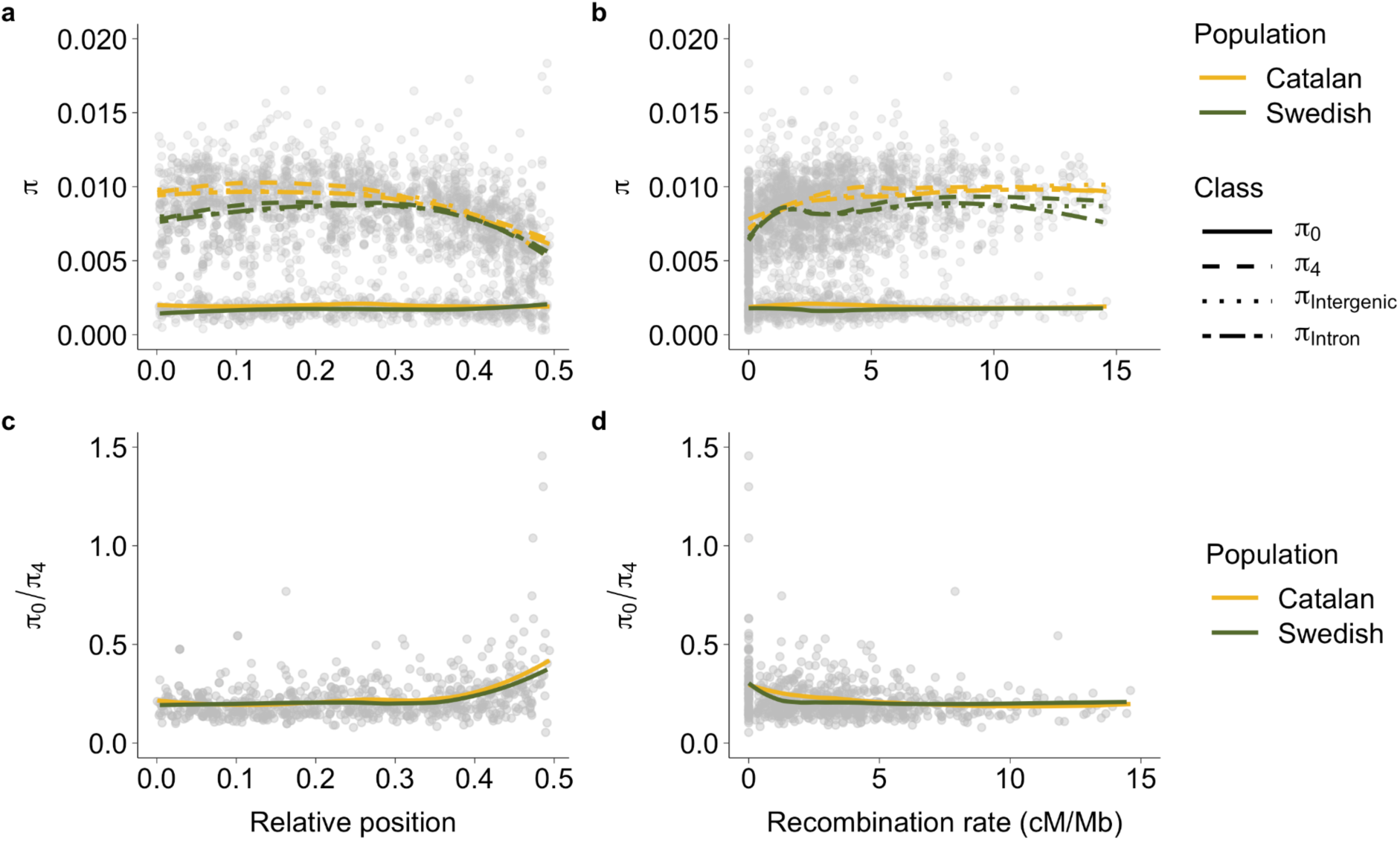
a) Regional distribution of nucleotide diversity (*π*) in zero-fold (*π*_0_), four-fold degenerate (*π*_4_), intergenic and intronic sites. b) Nucleotide diversity (*π*) as a function of the recombination rate in each population estimated in 2 Mb windows. c) Zero-fold/four-fold diversity (*π*_0_/ *π*_4_) along the chromosome (relative position from chromosome center (0) to chromosome end (0.5)) for the Swedish (green) and Catalan (yellow) *L. sinapis* populations. d) Zero-fold/four-fold diversity (*π*_0_/ *π*_4_) as a function of the recombination rate in each population estimated in 2 Mb windows. Lines represent local regression lines (LOESS).

**Table 2.** Genome wide estimates of pairwise nucleotide diversity (*π*) for different nucleotide site categories and the ratio of *π* at zero-and four-fold degenerate sites (*π*_0_ / *π*_4_) for each population. The ranges for chromosome specific estimates are given in parentheses. The statistics (W) and the p-values correspond to Wilcoxon rank sum tests with continuity correction.

| Sites | Catalan (*10<sup>-2</sup>) | Swedish (*10<sup>-2</sup>) | W | p-value |
| --- | --- | --- | --- | --- |
| $\pi_0$ | 0.19 (0.12 – 0.28) | 0.16 (0.09 - 0.19) | 1,183 | 2.43*10<sup>-05</sup> |
| $\pi_4$ | 0.92 (0.51 – 1.26) | 0.82 (0.43 – 1.02) | 1,124 | 2.73*10<sup>-04</sup> |
| $\pi_{\text{Intergenic}}$ | 0.90 (0.59 – 1.19) | 0.80 (0.50 - 0.97) | 1,258 | 7.05*10<sup>-07</sup> |
| $\pi_{\text{Intron}}$ | 0.87 (0.57 – 1.10) | 0.79 (0.48 - 0.96) | 1,175 | 3.44*10<sup>-05</sup> |
| $\pi_0 / \pi_4$ | 20.37 (13.97 – 34.85) | 19.54 (16.56 – 31.40) | 900 | 1.52*10<sup>-01</sup> |

We applied linear models to investigate the association between different diversity estimates and the variation in the regional recombination rate in each population in more detail. These analyses revealed that window-based estimates of diversity at all site categories, except *π*_0_ (R^2^ < 0.01, p-value > 2*10^-01^), were significantly associated with the recombination rate in both populations (R^2^ = 0.05-0.13, permuted p-values < 1*10^-03^; Fig. 3, Suppl. Fig. 4). Consequently, there was a negative relationship between the recombination rate and *π*_0_ / *π*_4_ in both the Catalan (R^2^ = 0.04, p-values < 1*10^-03^) and the Swedish population (R^2^ = 0.05, p-values < 1*10^-03^), suggesting that the overall recombination landscape has been stable enough to have a similar effect on diversity and efficacy of selection in both populations since they diverged (Fig. 3, Suppl. Fig. 4). The relationship between recombination rate was non-linear with a stronger association in low recombination regions, indicating a limited effect on diversity and efficacy of selection when the recombination rate increases above a certain level (∼ 2 cM / Mb).

Global and regional estimates of recombination and diversity could be affected by other covarying variables, such as different genomic features, nucleotide composition and demographic processes. In an attempt to account for these variables and investigate the relative effect of karyotype rearrangements, we quantified how observed changes in recombination rate between populations were associated with changes in diversity between the populations. This analysis was performed using homologous segments in ancestral chromosome blocks (Fig. 4). We focused on ancestral chromosomal units where the fission / fusion has been inferred to occur after the split from the closest outgroup species (*L. reali*), and cases where incomplete lineage sorting or reuse of chromosomal break-points have resulted in differences in karyotype between the two populations. This analysis unveiled a significant positive association between the change in recombination rate and intergenic *π* (Linear regression, R^2^ = 0.12, permuted p-value = 7.99*10^-03^; Fig. 4). There was also a positive relationship between recombination and intronic *π*, but this trend was not significant after correction for multiple testing. We found no associations between recombination rate change with changes in *π* for other site classes or with *π*_0_ / *π*_4_, possibly an effect of stronger selective constraints for those sites (Suppl. Fig. 5).

**Fig. 4.**
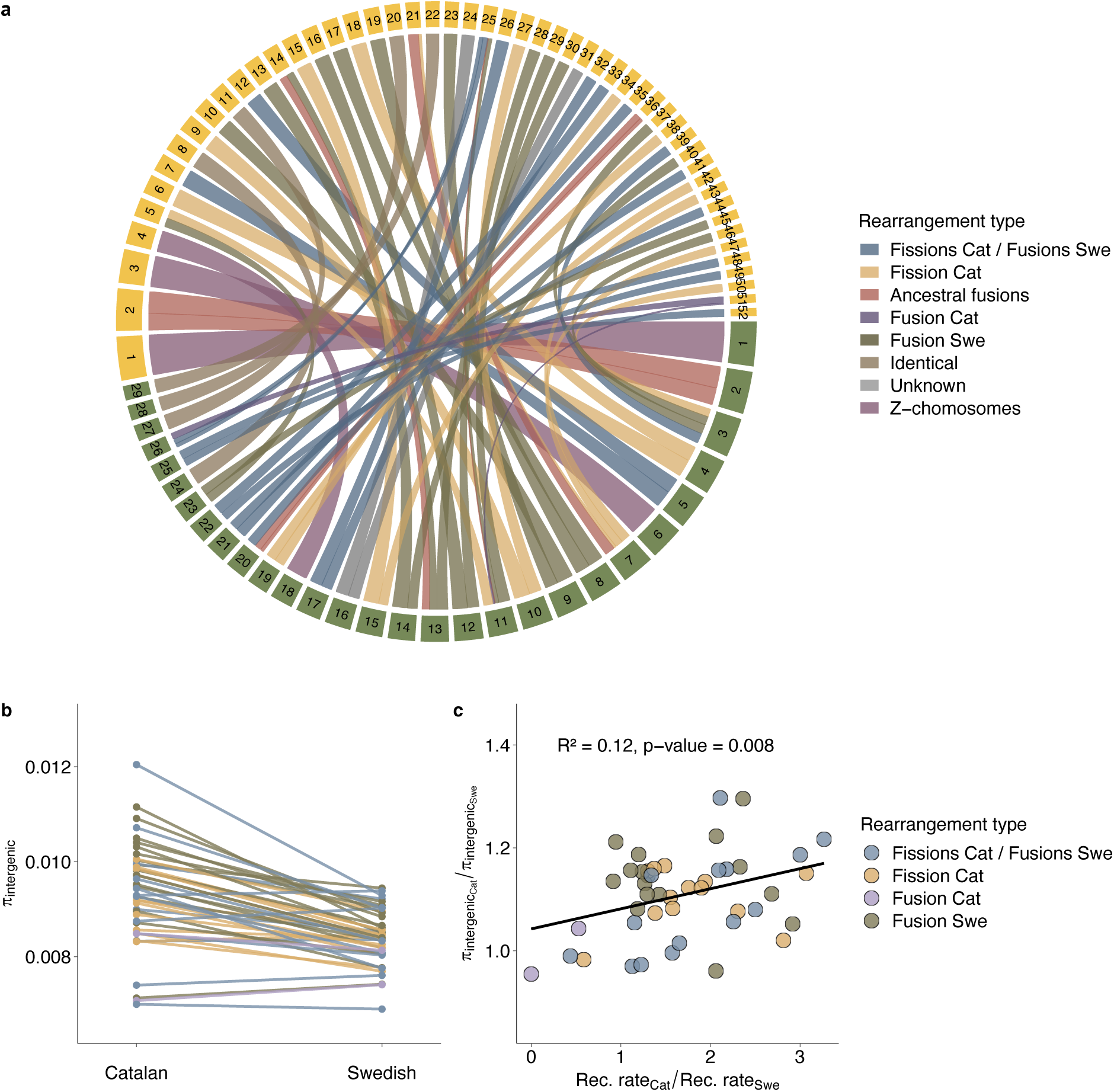
a) Chromosomal rearrangements between Swedish (green squares) and Catalan (yellow squares) population of *L. sinapis*, the bands representing homology are colored by fission/fusion history. Chromosomes are ordered by size in each respective population (data from [21]) b) Diversity estimates (*π*), specifically for intergenic sites across ancestral chromosomal units in each population. c) The ratio between Catalan and Swedish *π*, as a function of the ratio between Catalan and Swedish recombination rate for intergenic sites in each ancestral chromosomal unit with known fusion/fission history. The line represents the slope from a model II linear regression (R^2^ and permutated p-value on top).

To summarize, the analysis shows that chromosomal rearrangements have affected the recombination landscape and changes in the level of standing genetic variation in intergenic sequences in *L. sinapis*.

## Discussion

### General

Here we characterized the effects of extensive within-species chromosomal rearrangement on recombination and total pairwise reshuffling rate variation and combined this with analyzing the associated influences on levels of genetic diversity and the efficacy of natural selection. In summary, we found i) that karyotype rearrangements affect both the genome-wide and the regional recombination rate, ii) that the number of chromosomes and their relative size are major determinants of the recombination rate, but that karyotype changes have very limited effects on the total pairwise reshuffling rate, and iii) that recombination rate changes associated with chromosome fissions and fusions have detectable effects on the neutral diversity at intergenic sites, but not on the efficacy of selection.

### Recombination rate analysis

Our linkage map data revealed that the recombination rate was higher in the Catalan than in the Swedish population. This is expected given that chromosome size previously has been shown to be strongly associated with recombination rate in Lepidoptera [22,24]. We found that both *Leptidea* populations had global recombination rates in parity with genome-wide estimates in other Lepidoptera species, although direct comparison here is difficult since different methods have been applied [22,26,27]. However, the effective recombination rate per gene is likely higher in *L. sinapis*, since it has a large genome size (686 Mb) compared to many other butterfly species [28]. The difference in recombination rate between populations was lower than predicted from karyotype differences if the average chromosome map length would have been 50 cM. One potential explanation for this deviation is the occurrence of multiple crossovers in large chromosomes, an explanation which is supported by our observation that excess recombination (> 50 cM / chromosome) increases linearly with chromosome size. The presence of multiple crossovers per chromosome indicates that butterflies are not limited to one crossover per meiosis, as *C. elegans* [29], but rather that the number of crossovers is dependent on chromosome size like in other holocentric species [30]. Notably, we also found that smaller chromosomes sometimes showed a lower average crossover rate than expected if crossovers are necessary for correct segregation during meiosis in males (i.e. < 50 cM / chromosome). This could potentially be a result of very closely located double crossovers or single crossovers occurring close to the chromosome ends. Such events are likely largely undetected with our marker density. Another explanation, perhaps less likely, could be that correct segregation in male meiosis might not require chiasma formation at all, as has been observed in the achiasmatic Lepidoptera females. Unpaired univalents have for example been observed during meiosis in crosses between chromosome races of *L. sinapis* [31]. Another factor that likely can influence the difference in recombination rate between larger and smaller chromosomes is the difference in telomeric and subtelomeric proportions where the recombination rate was observed to be significantly reduced.

We also found that the recombination landscape along many of the larger chromosomes was bimodal, with a tendency towards lower recombination rate in the center. Similar patterns have been observed in other recombination analyses in Lepidoptera [22,24,32], but typically not as pronounced as in other organisms [15,29,33]. There are two potential explanations for this bimodality in recombination frequency along larger chromosomes. First, it is likely that interference will occur when > 1 chiasmata are formed on a single chromosome [34]. This would likely result in chiasma formation towards chromosome ends, but not all the way towards the telomeres where recombination seems to be inhibited [35]. An alternative, albeit not exclusive explanation, is that double-strand break initiation is directed away from the center and closer towards the telomere regions, as a consequence of physical proximity when the chromosome ends aggregate close to the nuclear membrane (the ‘meiotic bouquet’) during early meiotic stages [9].

### Reshuffling rate estimates

Changes in chromosome numbers as a consequence of fissions and fusions may have an effect on the reshuffling rate of genetic variants, since there are more opportunities for independent segregation with an increasing number of chromosomes. We therefore investigated the relative contribution of both independent segregation and recombination to the total reshuffling rate in the two *L. sinapis* populations. The probability of interchromosomal pairwise reshuffling was close to the maximum (50%) in both populations, which means that the difference in number of chromosome pairs between the Catalan and the Swedish *L. sinapis* (29 and 52, respectively) has a marginal effect on difference in total reshuffling rate (diminishing return when chromosome numbers exceed n = 25 - 30; [8]. Notably, most eukaryotes have a diploid chromosome number << 50, with a mean 2n = 17.05 [10] and reshuffling has been proposed to be beneficial for short term distribution of diversity in the offspring [36]. However, the relatively limited range of chromosome numbers in natural populations indicates that there are only minor selective advantages of increasing the chromosome number above a certain threshold. Possible explanations can be that beneficial or co-adapted allele combinations can be inherited together when there are fewer chromosomes and there could possibly be increased risks for segregation errors with higher chromosome numbers. Here we show that the reshuffling between chromosomes contributes much more (orders of magnitude) than intrachromosomal processes to the total reshuffling rate in both populations, confirming the paramount importance of independent assortment of chromosomes for distributing variation in the gametes. Homologous recombination can therefore be seen as a long-term mechanism for disruption of linkage (and Hill-Robertson-interference) between closely positioned loci within chromosomes. It has for example been shown that increasing the number of recombination events is more important than increasing the number of chromosomes for the efficiency of selection [37]. Although we found that the intrachromosomal reshuffling contributed marginally to the total rate, we also found that the pairwise reshuffling was almost twice as high in the Swedish population compared to the Catalan population. This is obviously an effect of chromosome size, analogous to the positive correlation between genetic map length and intra chromosomal reshuffling rate in some plants [15]. The underlying process must be that > 1 crossover can occur on larger chromosomes, and that these crossovers are located distantly enough to increase intrachromosomal reshuffling significantly (double crossovers increase the intrachromosomal pairwise reshuffling if they are far apart-two crossovers located at ¼ of the chromosome length from the terminal results in as high pairwise reshuffling as one crossover in the center). Our data suggest that recombination events are limited by interference mechanism that both influence the distance between, and the total numbers of, chiasmata occurring on a single chromosome. Interference seems to be ubiquitous among sexually reproducing organisms, but the precise mechanism is not described for many species [34,35]. We also found that chromosomes < 20 Mb roughly have a recombination distance of 50 cM, similar to the situation in *H. melpomene* where all chromosomes are < 20 Mb [26,38]. These results spur interest in further investigating the mechanistic basis of recombination interference in butterflies in general. In summary, we observed a higher recombination rate in the smaller chromosomes, but this was compensated for by a higher intrachromsomal reshuffling in larger chromosomes. Hence, we proceeded by assessing if these differences and similarities in recombination and reshuffling can affect levels of genetic diversity.

### Associations between recombination and genetic variation

To get more information about how chromosome rearrangements and recombination rate differences between populations and genomic regions are associated with genetic diversity, we estimated a set of population genetic summary statistics at different site categories. Our results show that there was a significantly higher overall genetic diversity in the Catalan population than in the Swedish population. Such a difference could obviously be a consequence of differences in demographic history or other evolutionary processes that affect the populations differently. First, we know that *L. sinapis* populations inhabiting warmer climatic regions (e.g. Catalonia) are multivoltine with 3-4 generations per year, while the populations at higher latitudes (e.g. Sweden) predominantly are univoltine [39]. A shorter generation time leads to a higher input of novel mutations per time unit, but this should only affect the segregating neutral diversity in the populations differently if they are not at mutation-drift-equilibrium [40]. Second, the demographic histories are likely different between the two populations. While Catalonia likely represents a refugium from the last glaciation period, Sweden must have been colonized well after the ice sheet retracted. This founder event in the north likely occurred with influx of individuals from another refugial population, potentially as far east as central Asia [32,41]. Hence, both the shorter generation time and the absence of obvious founder events could have led to a higher genome-wide level of genetic diversity in the Catalan population.

To further investigate the effects of recombination on patterns of genetic diversity we therefore focused on intra-genomic variation in each population separately. We found that the levels of genetic diversity were heterogeneous across chromosomes and there was no association between genetic diversity and chromosome size after excluding the Z-chromosomes from the analysis. The latter result could be a consequence of the dynamic ongoing karyotype rearrangements in this species, and fits well with linkage-disequilibrium-based recombination rate estimates in the Swedish population [32]. The recurrent fissions and fusions that must have occurred over the divergence time of the populations should mean that both the size of chromosomes and (consequently) the recombination rate has been under constant change, which would reduce the association between current chromosome states, recombination rates and levels of genetic diversity. In addition, a time lag between changes in recombination rate and subsequent changes in genetic diversity is expected, which should dilute the signals further [42]. However, we did actually detect a significant genome-wide association between genetic diversity and recombination rate in both populations. This suggests that the split between population occurred a sufficiently long time ago for recombination rate differences to affect genetic diversity. Here, the association was strong at the lower end of the recombination rate spectrum and reached a threshold at moderate recombination rates, with similar levels to those previously detected in *Drosophila* [43]. This is also in line with theoretical predictions of the association between recombination and genetic variation [44].

The regional distribution of genetic diversity along the chromosomes were in line with the expectations [3], following the trajectory of the recombination landscape in both populations. It should be noted that the chromosomal rearrangements in *L. sinapis* mainly have involved fusions and/or fissions of comparatively large chromosome parts [21]. The distribution of the recombination landscape on chromosomes could hence be partly maintained even after fusion of two smaller chromosomes or a fission of a larger chromosome -i.e. unimodality in small chromosomes with recombination predominantly occurring at the center and bimodality in larger chromosomes with recombination events towards, but not at, chromosome ends. However, this relative stability obviously depends on the history of the merged chromosome units. Recent fusions, for example, would be expected to have comparatively high genetic diversity as a result of the higher recombination rate on the previously smaller chromosomes, while more ancient fusions would have a lower level of genetic diversity as a consequence of a more long-term recombination rate reduction [25]. It should be noted that fission / fusion polymorphisms are known to be segregating in the populations [21] and that the linkage maps used here are composite maps of different families which may add some uncertainty to the estimates of recombination rates. To further investigate the extent of potential individual differences in recombination rate as a consequence of karyotype variation between families within a specific population, we would need to produce individual recombination maps for males homozygous for each karyotype.

Both cytogenetics [20], linkage mapping and genome assemblies [21] have revealed the existence of an unusual sex-chromosome system in the *Leptidea* genus where e.g. *L. sinapis* has a karyotype including three Z-chromosomes and three W-chromosomes. Of the three Z-chromosomes, one is homologous to the inferred ancestral Z-chromosome (Z1) in Lepidoptera while the other two (Z2, Z3) seem to have been recruited as sex-chromosomes in the *Leptidea* genus specifically [20]. To understand how this relatively new evolution of sex-chromosomes have affected the recombination and diversity landscapes, we compared the different Z-chromosomes to the set of autosomes. The results from this analysis showed that the Z-chromosomes had a lower neutral genetic diversity than autosomes. This is expected given that the *N_e_* of a Z-chromosome should be approximately ¾ of any autosome, under the assumption of equal sex ratios. A reduced level of genetic diversity on the Z-chromosome has also been observed in the *Heliconius melpomene* and the Monarch butterfly, but not in the moth *Manduca sexta* [45,46], suggesting that relative *N_e_* of Z-chromosomes and autosomes might vary between species, or that other factors than mutation-drift balance drive relative levels of genetic diversity between chromosome classes. An interesting result was that while Z1 and Z3 had similar *π*_0_ / *π*_4_-ratios as the autosomes, Z2 had a dramatically higher *π*_0_ / *π*_4_-ratio compared to the other chromosomes in both populations. The higher ratio was a consequence of an elevated *π*_0_, indicating reduced selective constraints on Z2 (or possibly a group of genes under strong positive selection). This result is somewhat puzzling since Z2 is the most structurally conserved of all chromosomes within *Leptidea* [21,47] and the observation, although outside of the scope of this article, definitely merits further investigations into the evolutionary forces underlying sex-chromosome formation and maintenance in this genus.

The efficacy of selection translates to both removal of deleterious mutations and fixation rate of adaptive mutations. Hence, under the assumption that most novel mutations are neutral or slightly deleterious, i.e. the nearly neutral theory [48], we expect an increase in the *π*_0_ / *π*_4_-ratio with reduced efficiency of selection. Since the efficiency of selection is dependent on *N*_e_, the frequency of (slightly) deleterious mutations is expected to segregate at higher rates in regions of low recombination [49]. Our data showed a negative association between the recombination rate and the *π*_0_ / *π*_4_-ratio in both populations, an observation supporting the nearly neutral theory. However, we did not detect any significant difference between the two populations, which would be expected given the difference in overall recombination rate (i.e. a lower *N*_e_ in the Swedish population). In addition, there was no detectable effect of the variance in recombination rate in rearranged chromosomes on the ratio of *π*_0_ / *π*_4_. This indicates that the average recombination rate is high enough to efficiently remove slightly deleterious mutations in both populations, consistent with the observations that selection efficiency can be maintained at a stable level even at relatively low recombination frequencies [50]. In the genome-wide analysis we observed that the *π*_0_ / *π*_4_-ratio approached the asymptote when the recombination rate was > 2 cM / Mb. Hence, an increase in the recombination rate above that level does not seem to affect the efficiency of selection in this system. It is also possible that the relatively short divergence time between these populations [51], in combination with the extensive chromosome changes, affects our power to detect signals of selection.

We found a positive association between the variation in intergenic diversity change and the variation in recombination rate change across ancestry blocks, with recombination rate explaining > 12 % of the variation in intergenic diversity in rearranged chromosomes. A positive association between recombination rate and genetic diversity has been observed in many different taxa [52] and our data lends further support for that recombination is a more potent mechanism than pairwise chromosome reshuffling for breaking linkage disequilibrium and, hence, maintenance of genetic diversity [37]. It is noteworthy that we did not see any significant association between changes in recombination rate in rearranged regions and genetic diversity at other site categories (introns, *π*_0_, *π*_4_). This is likely a consequence of temporal differences in the effects of reduced linkage disequilibrium on genetic diversity at sites under direct and indirect selection pressure.

## Conclusion

We investigated the difference in recombination rate between two populations of *L. sinapis* with extreme difference in karyotype structure and chromosome count. We show that karyotype evolution directly affects the global recombination rate, but that recombination has a limited effect on total pairwise reshuffling, partly as a consequence of multiple crossovers in larger chromosomes. A key observation from our data was a significant difference in neutral genetic diversity (but not on the efficacy of selection) in regions where chromosome rearrangements have affected the recombination rate. Finally, we validate the importance of recombination rate variation driving differences in maintenance of nucleotide diversity between populations at an early stage of divergence.

## Methods

### Linkage map

We used Swedish (n = 6 families; 186 offspring) and Catalan (n = 6 families; 186 offspring) full-sib families of *L. sinapis* to develop RAD-seq based linkage maps for two populations that represent the most extreme karyotype variants in the species. Details about family structure, RADseq data generation and preprocessing are available in [21]. To get recombination rate information, the linkage map markers were anchored on recently established chromosome-level assemblies for each respective population [21]. The filtered reads were mapped to the genome assembly from each population using bwa *mem* [53] with default options. The mapped reads were sorted with samtools *sort* and filtered based on quality (samtools *view-q 10*), and reads with multiple mapping locations were removed with a custom script [54]. Mapping coverage was assessed with Qualimap [55] and only individuals with > 100,000 mapping reads were included in the study, resulting in 184 and 178 offspring in the Swedish and Catalan pedigrees, respectively. Samtools *mpileup* [56] with options minimum mapping quality (*-q*) 10 and minimum base quality (*-Q*) 10 was used to identify variants, which were converted to genotype likelihoods using *Pileup2Likelihoods* in LepMap3 with default settings [57]; minimum coverage = 3 per individual (*minCoverage* = 3) and < 30% of the individuals allowed to have lower coverage than minimum coverage (*numLowerCoverage* = 0.3). Only markers mapping to chromosome-sized scaffolds in each respective population were retained for downstream analysis. LepMap3 [57] was used to construct linkage maps for the two populations separately applying the following steps. Informative markers were identified with *ParentCall2* using default settings, except for *ZLimit* which was set to 2 to identify markers segregating as sex chromosomes. Non-informative markers were removed with *removeNonInformative* = 1. Markers showing segregation distortion, minimum allele frequency below 0.05 or absence in more than 50% of the individuals were excluded with *Filtering2* (options *dataTolerance* = 0.00001, *MAFLimit* = 0.05, *missingLimit* = 0.5). In addition, only markers represented in at least four families in each population, were retained (*familyInformativeLimit* = 4). As a first mapping step, markers were assigned to linkage groups with *SeparateChromosomes2* using female informative markers only (*informativeMask* = 2). The optimal LOD-score threshold was empirically estimated to 11 for the Swedish and 7 for the Catalan population. Markers that were informative in either males or both sexes were subsequently added with *JoinSingles2All* with the same LOD-score thresholds. Linkage groups covering several scaffolds were split based on information from the physical assembly before ordering the markers along the chromosomes with the module *Ordermarkers2*. Only male informative markers (*informativeMask* = 1) were used for the ordering since females are achiasmatic (*recombination2* = 0). We set a penalty for diverting from the order in the genome assembly with likelihood chain (*usePhysical* = 1 0.1) and *improveOrder* = 0. The ordered linkage maps were visually inspected in R and uninformative markers (extending the map ends) were removed. After manual removal of uninformative markers, *OrderMarkers2* was run again with the same settings (see above) and the final genetic distances were calculated using Kosambi’s map function to account for multiple recombination events along the chromosome.

### Recombination estimation

Regional recombination rates were estimated by dividing the genetic distance with the physical interval between each marker pair. Weighted means for 2 Mb windows, per chromosome and per population were obtained by weighting the recombination rate at each interval by the proportion of the physical distance. Interpopulation comparisons of the weighted mean recombination rates were performed with Wilcoxon Signed rank test and potential associations between recombination rate and chromosome length were assessed with Spearman’s rank correlation coefficient, as implemented in cor.test in R [58]. To compare the observed recombination rate with the expected recombination rate for exactly one or two crossovers per chromosome per meiosis, we calculated the expected rates as 50 cM / chromosome length and 100 cM / chromosome length, respectively.

### Pairwise reshuffling

The probability of total pairwise reshuffling is the probability that two random loci are originating from different parental alleles during gamete formation. The total pairwise reshuffling has two components; the interchromosomal component describing the effects of independent assortment and the intrachromosomal component describing the effect of recombination. The variation in recombination rate along each chromosome is also corrected for. The probability of interchromosomal pairwise reshuffling was estimated by the probability of two loci being located on different chromosomes with independent assortment of the chromosomes during meiosis (1 – *sum* ((*chromosome length* / *assembly lenght*)^2^)) ∗ 0.5 [8]. The intrachromosomal probability of pairwise reshuffling per chromosome was inferred using the genetic and physical distances from the linkage maps. 1,000 pseudomarkers were placed along each chromosome and the number of pairwise differences in origin was calculated based on the probability of recombination between each pair of pseudomarkers [8]. The sum of pairwise differences was divided by the total number of comparisons for each chromosome. The intrachromosomal contribution to the total reshuffling rate was estimated by summing up the contribution of each chromosome, weighted by the probability that two loci are located on the same chromosome (chromosome length / assembly length)^2^. The total pairwise reshuffling was calculated as the sum of the interchromosomal and intrachromsomal contribution [8].

### Population genetic summary statistics

Previously available resequencing data from 10 Swedish and 10 Catalan individuals were used for population genetic analysis (Talla et al., 2019). Positions with Phred-score < 33, sequencing adapters and seven bases from the 3’-ends of all reads were trimmed using TrimGalore 0.6.1 (https://github.com/FelixKrueger/TrimGalore), a wrapper for cutadapt 3.1 [59]. Trimmed reads were mapped to the genome assemblies for each respective population [21] using bwa *mem* [53]. Variant calling was performed using GATK 4.2.0.0 [60]. Base quality score recalibration was performed using an initial variant calling round, keeping a set of variants with good support as recommended by GATK [61]. Subsequently, a second variant calling round was performed on recalibrated mapped reads. A final filtering step of all sites (variant and invariant sites) was applied using BCFtools *filter* 1.16-1 [62] to mask genotypes overlapping annotated repeats and genotypes with a coverage < 5 or > 25. The applied options for these filtering steps were *--mask-file $REPEAT_GFF, --soft-filter “REPEAT”,* and *-i FMT/DP > 5 & FMT/DP < 25’ --set-GTs*.

Nucleotide diversity (*π*) for intronic, intergenic, 0-fold and 4-fold sites were estimated in 50 kb windows with the software pixy 1.2.5.beta [63]. Coordinates for 0-fold and 4-fold sites were obtained with a custom script (modified from https://github.com/simonhmartin/genomics_general) and genomic features (intron, intergenic) were extracted using BEDTools 2.29.2 *complement* [64]. Potential associations between *π* and chromosome length were assessed with Spearman’s rank correlation, as implemented in cor.test in R [58]. We used the ratio of *π*_0_ / *π*_4_ as a proxy for selection, reflecting the efficacy in removal of deleterious mutations, since most protein changing mutations are either deleterious or neutral according to the nearly neutral theory [48].

Potential associations between the recombination rate and *π* were assessed by binning data into 2 Mb genomic windows for each population separately. Population genetic summary statistics were also estimated in ancestry blocks (ancestral chromosomal units), i.e. previously identified homologous regions of chromosomes in which no rearrangements have occurred between the populations [21]. We included ancestral chromosomal units involved in fissions/fusions after the split from the closest sister species (*Leptidea reali*), or where incomplete lineage sorting or reuse of breakpoints have resulted in a fission in one population or a fusion in the other. To investigate if observed changes in recombination rate between populations has resulted in a corresponding change in *π*, we compared the ratio of the Catalan and the Swedish *π* with the observed recombination rate ratio between the populations within each ancestral chromosomal unit. Both the recombination rate and the *π* estimates are random variables, so we applied a model II linear regression using ordinary least squares (OLS) and a permutation test with 1,000 permutations to determine the significance of the slopes, as implemented in the R-package lmodel2 [65].

## Acknowledgements

This work was funded by the Swedish Research Council (VR research grant #019-04791 to N.B.). The authors acknowledge support from the National Genomics Infrastructure in Stockholm funded by Science for Life Laboratory, the Knut and Alice Wallenberg Foundation and the Swedish Research Council, and SNIC/Uppsala Multidisciplinary Center for Advanced Computational Science for assistance with massively parallel sequencing and access to the UPPMAX computational infrastructure. The project was also supported by NBIS/SciLifeLab long-term bioinformatics support (WABI).

## Data availability statement

The data used in this article are from public repositories. The genome and linkage map data for *L. sinapis* are available at the European Nucleotide Archive (ENA) under accession number: PRJEB58697 (genome assemblies) and PRJEB58905 (pedigree RADseq data). The population data are available at ENA under accession number: PRJEB21838. All in-house developed scrips are available on GitHub (https://github.com/EBC-butterfly-genomics-team).

## Supplementary information

Link to supplementary information document (Supplementary Information provided as a separate file).

## Notes

### Competing Interest Statement

The authors have declared no competing interest.

